# Potential Outbreak in Brazil by Equine Herpesvirus Type 2 (EHV-2): Detection, Clinical and Genetic Analysis of Equine Gammaherpesvirus 2

**DOI:** 10.1101/809053

**Authors:** Sofia Cicolo, Angélica Cristine de Almeida Campos, Cairo Monteiro de Oliveira, Thais Poltronieri, Luiz Gustavo Bentim Góes, Vanessa Barbosa da Silveira, Carla Meneguin Barbosa, Raquel Yvonne Arantes Baccarin, Danielle Bruna Leal Oliveira, Carla Bargi Belli, Edison Luiz Durigon

**Affiliations:** Department of Veterinary Surgery. Faculdade de Medicina Veterinária e Zootecnia da Universidade de São Paulo, São Paulo, SP, Brazil; Department of Microbiology. Instituto de Ciências Biomédicas da Universidade de São Paulo, São Paulo, SP, Brazil; Department of Veterinary Internal Medicine. Faculdade de Medicina Veterinária e Zootecnia da Universidade de São Paulo, São Paulo, SP, Brazil

**Keywords:** Horse, foal, herpesvirus, pneumonia, neurological signs, abortion, EHV-2

## Abstract

Equine herpesvirus (EHV) commonly affects horses causing neurologic and respiratory symptoms beside spontaneous abortions, meaning huge economic losses for equine industry worldwide. In foals the virus can facilitate secondary infections by *Rhodococcus equi*, important in morbidity and mortality in equines. A total of five genotypes of EHV were previously described in Brazil including EHV-1, EHV-2, EHV-3, EHV-4 and EHV-5. EHV-2 genotype had only been previously described in Brazil in asymptomatic animals. We report the dead of 11 foals in Middle-west region of Brazil showing respiratory and neurological symptoms, as well as several abortions in mares from the same farm. Laboratory exams revealed neutrophilia leukocytosis. Necroscopic and histopathologic findings were suppurative bronchopneumonia and ulcerative enteritis. Molecular assays point to the absence of the bacteria *Rhodococcus equi* and other EHV. The presence of EHV-2 DNA was confirmed by sequencing in serum sample from one foal. This is the first confirmed outbreak of EHV-2 causing disease in Brazilian horses with confirmed presence of the virus. EHV-2 has been neglected and could be highlight the important role of EHV-2 in equine respiratory disease and spontaneous abortions in equid in Brazil.

**Author Summary:** Equid herpesvirus type 2 (EHV-2), a gammaherpesvirus, has been neglected and can cause respiratory infection, abortion and in extreme cases neurologic symptoms and dead.

Outbreaks of EHV-2 never was described in Brazil and can be critical to losses in equid industry. Our report provides evidence for a direct link between the EHV-2 strain and the outbreak in foals. We proved the clinical, histopathological and phylogenetic analysis of segment of EHV-2 polymerase.

In summary, our study confirm that Gammaherpesvirus 2 is circulating in Brazil and the EHV-2 is a cause of the first outbreak in foals showing the important role of EHV-2 in equid respiratory disease.

## 1. Introduction

Horses are the natural host of five distinct equine herpesvirus (EHV), members of the Herpesviridae family (Order Herpesvirales) distributed in two groups Alphaherpesvirus and Gammaherpesvirus (1). EHV-1, EHV-3, EHV-4, EHV-8 and EHV-9 are members of the Alphaherpesvirinae subfamily, genus Varicellovirus and EHV-2, EHV-5 and EHV-7 are classified in the Gammaherpesvirinae subfamily in the Percavirus genus (EHV-2 and EHV-5) and EHV-7 in the Rhadinovirus genus (ICTV 2018).

Equine herpesvirus type-2 (EHV-2) are known as the cause of acute infections and long latency in horses. This virus has been frequently correlated to immunosuppression in foals, upper respiratory tract and chronic pulmonary disease, keratoconjunctivitis, general malaise, abortion (2), poor performance and granulomatous dermatitis (3,4).

EHV-2 infection usually occurs soon after birth (5). However, the detection of EHV-2 DNA in the lungs of the aborted fetus indicate that this virus may be capable to cross the placental barrier and infect the foal in gestational period (6). Beside the absence of an effective vaccine against EHV-2, Belák et al. (7) showed that producing hyperimmune serum of a local strain of EHV-2 and vaccinating foals with repeated applications may prevent the disease.

Apparently, EHV-2 disease presents a two-phase evolution in young foals. The initial phase begins with an occasional appearance of nasal discharge and slight increase in body temperature fallowed by a temporary recover and the development of a severe respiratory disease, pneumonia, pulmonary abscesses, high fever and death. The second phase in characterized by complications aggravated by the bacteria *Rhodococcus equi* with the invasion of the respiratory tract (7,8). This secondary infection causes bronchopneumonia, intestinal manifestations, septic arthritis and osteomyelitis, uveitis and sepsis jeopardizing the foal athletic career or even leading to death (9,10).

EHV-2 are endemic is horses population worldwide been identified in horses without clinical symptoms or presenting respiratory disease (4,11–13). In South America a study conducted in Argentina report the circulation of EHV-2 in thoroughbred horses presenting respiratory symptoms also indicating a high prevalence in this equine population (11). The presence of EHV-2 in Brazilian horse herd was only recently confirmed by the detection of herpesvirus DNA in nasal swabs of asymptomatics adult horses (12).

Despite the link between EHV-2 and acute infections, abortions and the possible link with poor performance in horses, the potential relation of EHV-2 in the development of clinical diseases in horses from Brazil were never analyzed. Here, we report the potential relation of EHV-2 on an outbreak causing the dead of 11 foals from Middle-West region of Brazil.

## 2. Material and Methods

### 2.1 Ethics Statement

The process was referred to the “Ethics Committee on Animal Use” from Biomedical Sciences Institute - University of Sao Paulo (CEUA-ICB-USP). This study was carried out in strict accordance with international recommendations. All efforts were made to minimize animal suffering.

### 2.2 Case Description

From the 14 foals that were born in a breeding farm from Middle-west region of Brazil during the year of 2018, 11 foals from “Mangalarga Marchador” breed (with age raging between 1 and 3 months) present respiratory symptoms such as dyspnea and pulmonary edema evolving to neurological signs (circling walking, lost look) and flexural limb deformities leading to death. Besides, four mares in the same farm and the same period had abortions.

One of the foals showing respiratory and neurological symptoms, a Brazilian saddle horse male (86 days age) was accompanied during this study. From this foal, clinical exam showed heart rate 104 bpm, respiratory rate 42 mpm, pale oral mucosa, capillary refill time 3 seconds, estimate dehydration 8%, reduced gut motility in all tract, rectal temperature 39.8 °C. Furthermore, the foal showed mild to marked dyspnea, nasal serobolous secretion, entropion in both eyes, conjunctival hyperemia, miosis, corneal opacity, presence of hypopyon and fibrin on eye anterior chamber. The complete blood count (CBC) disclosed leukocytosis (18100/mm^3^) by neutrophilia (12100/mm^3^) and hyperfibrinogenemia (800mg/dL).

Ceftiofur 10 mg/Kg, amikacin 20mg/Kg and flunixin meglumine 0.25mg/Kg IV were administered. 1% prednisolone acetate eye drops and 0.4% ketorolac tromethamine drops, one drop in each eye, every four hours was administered. Ringer lactato was the choice for IV fluids 5 mL/Kg/H, after two hours treating the patient died.

### 2.3 Sample Collection, Necropsy and Histopathology

Necropsy was performed following biosecurity standards procedures. Samples of serum, total blood, liquor and tissue from brain, gut, liver and spleen were collected and sent to histopathological analysis and pathogen molecular screening assays.

Serum, total blood, liquor and brain, gut, liver and spleen from one male (86 days age) Brazilian saddle horse breed “Mangalarga Marchador” were collected and sent to Clinical and Molecular Virology Lab ICB-USP (LabVCM).

### 2.4 Molecular assays

Quantitative Polymerase Chain Reaction (qPCR) assay in the detection of virulent *Rhodococcus equi* and EHV-1 and EHV-4 were performed in commercial laboratory.

Samples were transported in cold-chain logistic to the Laboratory of Clinical and Molecular Virology (LabVCM) from University of São Paulo. Total nucleic acids were extracted using NucliSENS® easyMAG® automatized platform (BioMerieux) following manufacturer’s instructions.

The polymerase herpesvirus genetic material was amplified by PCR and Nested-PCR assay targeting highly conserved region between all Herpesvirus subfamilies and capable to detect all species of EHV as described by (14) The positive sample was sent to Sanger Sequencing. Samples were also analyzed for the presence of distinct Flavivirus and Alphavirus by Pan-Flavivirus (15) and Pan-Alphavirus (16) Real-time SybrGreen qPCR assays.

The nucleotide sequences obtained were aligned using Geneious software (version Prime). Phylogenetic analyses were performed using MEGA7 software (version 7.0.2) using Maximum Likelihood (ML) with Neighbor-Joining (NJ) and GTR+◻+I model.

## 3. Results

Histopathological evaluation of lung tissue indicates Bronchi and bronchioles with hyaline membranes in their lumen. These findings are consistent with an acute, multifocal interstitial pneumonia, a marked coalescent associated with multifocal atelectasis. Bronchi and bronchioles have hyaline membranes in their lumen. In less affected areas there is a source of eosinophilic fibrillar material, possibly associated with edema, in an alveolar space and slight alveolar wall thickening. Multifaceted areas of marked alveolar wall rupture are observed in apical regions. Distinct tissues obtained from one of the 11 dead foal revealed no morphological changes in liver, spleen, kidney and heart.

During necropsy exam serobullous content was observed in the tracheal lumen, consistent with pulmonary edema; the lung was congested with areas suggestive of hepatization, in addition was observed to circular and whitish lesions of approximately one centimeter; engorgement of the mesenteric vessels, and cecal mucosa with marked hyperemia and diffuse hemorrhage (**Figure 1**).

**Figure 1:**
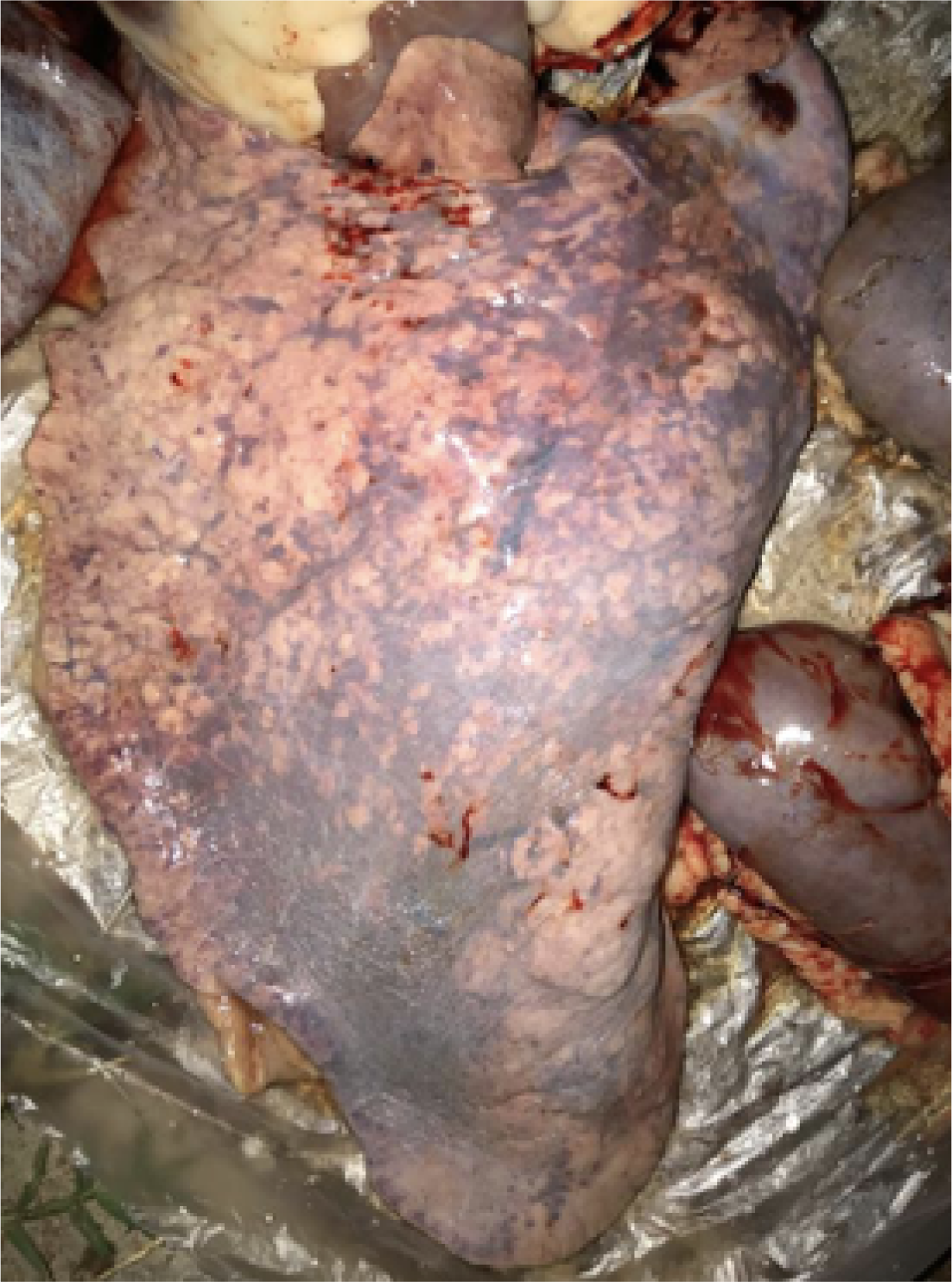
Necropsy exam. Foal lung presenting bronchopneumonia during necropsy exam.

Real-time quantitative polymerase chain reaction (qPCR) assays targeting virulent *Rhodococcus equi* and EHV-1 and EHV-4 were negative for all samples tested.

DNA from EHV-2 was detected in serum sample collected from one of the 11 the dead foal component of this study. All other tissues analyzed were negative n the same test. Serum and tissue were also negative for the presence of Flavivirus and Alphavirus RNA.

Serum shown amplification of 315 bp fragment in PCR-nested reaction and the presence of Gammaherpesvirus 2 was confirmed by sequencing and phylogenetic analysis (**Figure 2**).

**Figure 2:**
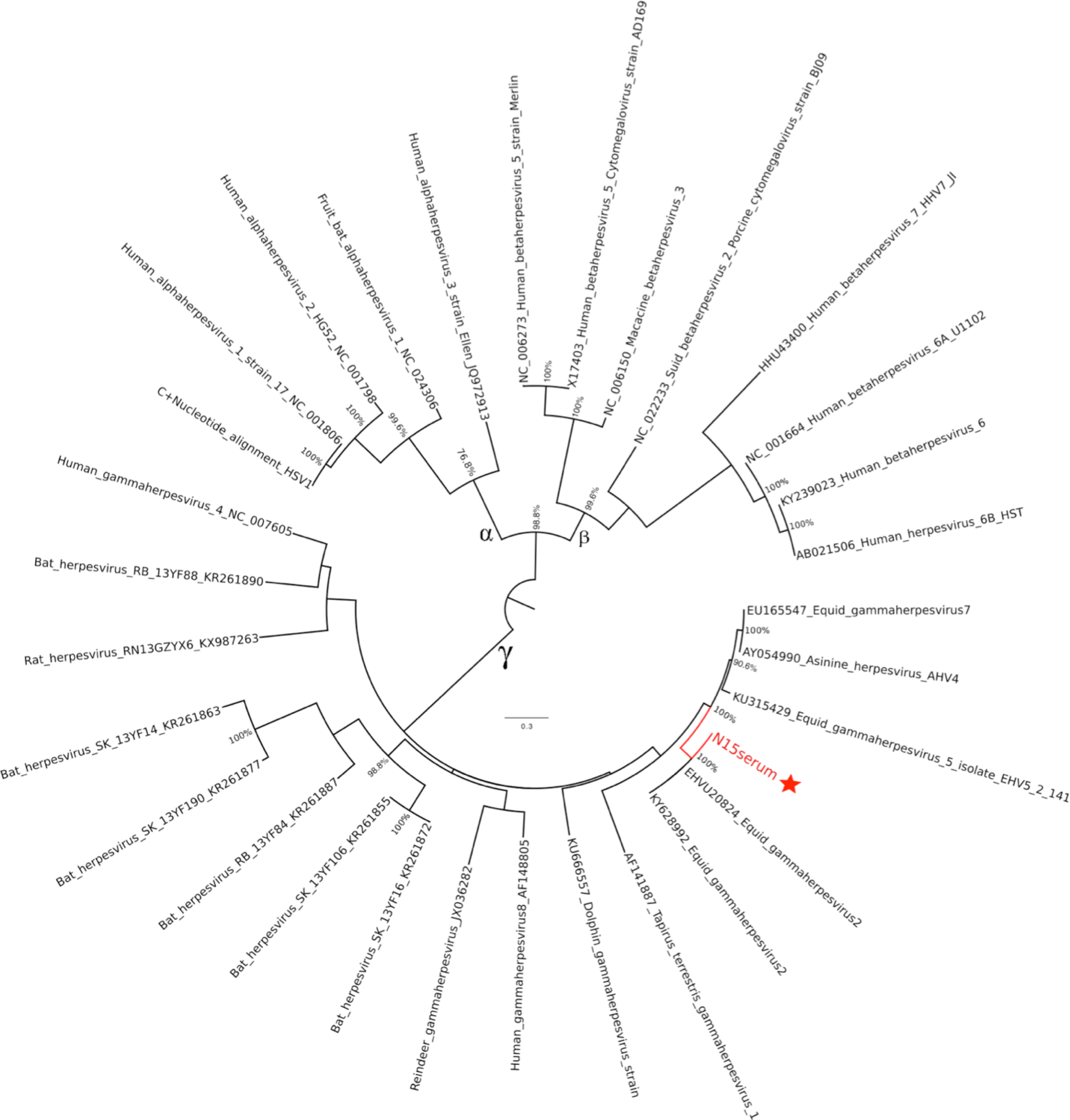
Maximum Likelihood Phylogenetic Tree. Tree reconstructed using a MEGA7 software with heuristic search and “GTR+gamma+I” algorithm. Bootstrap (10,000 bootstrap replicates) values greater than 50% are depicted in the figure and visualized in FigTree software (version 1.4.4). The analyzed sample are highlight in red segregating in Gammaherpesvirus clade.

## 4. Discussion

In this study we describe at first time the detection of EHV-2 in serum from equines presenting clinical symptoms in Brazil. In this mysterious outbreak disease related with death and abortions the epidemiologic data indicate the role of EHV-2 once the virus was detected in one serum sample.

The necropsy of the 11 horses indicate the same characteristics and cause of death. Samples related to one of this 11 death were collected and tested for distinct potential etiologic agents including *Rhodococcus equi* bacteria, Herpesvirus, Flavivirus and Alphavirus. The unique virus detected was EHV-2.

The genetic analysis confirm that sequences segregated in Equine Herpesvirus 2 clade and distance analysis revealed homogeneity (100% bootstrap values) with Equine Gammaherpesvirus 2 – EHV-2 (U20824 e KY628992) and shown 16.4% of distance with Equine Herpesvirus 5 (KU315429) and 14.5% distance of Equine Gammaherpesvirus 7 (EU165547). The obtained sequence was compared with herpesviruses from different hosts (equines, humans, monkeys, dolphins, bats, rodents and other mammals) representatives of the three main herpesviruses lineages (alpha, beta and gamma) available in GenBank databases.

EHV-2 presence in South America was previously reported in horses presenting respiratory clinical symptoms in Argentina (11) and only recently in health horses randomly sampled in Brazil (12). Until our data the relation of EHV-2 with horse diseases and abortion was reported only in Italy (17) and the role of EHV-2 in abortion still unclear (18).

However, Marenzoni et al. (19) detected EHV-1 and EHV-5 in 14% (9 animals) with different reproductive problems, including one co-infection with EHV-1 and EHV-2. The same authors detected the presence of EHV-5 in non-pregnant uterus, implying that it may have a role in reproductive dysfunction and have a negative consequence on the pregnant uterus.

EHV-2 can immunosuppress foals and allowed other bacterial infection and death. This represents an important economic, and affective lost for horse owners worldwide (20). In Brazil (21), we have a high prevalence of positive farms for *Rhodococcus equi*, a bacteria that causes several disease in foals and can have infectivity and pathogenesis increased by the presence of EHV-2.

EHV-1 is well known as a cause of abortion in mares (22), however EHV-2 transplacental transmission was demonstrated previously (6). The present study shows that not only EHV-1 but also EHV-2 can be an important cause of abortion in mares. In this report several mares aborted, even been vaccinated with commercial vaccines against equine herpesvirus. It demonstrates the necessity of a correct diagnostic in abortion cases and an effective vaccine against EHV-2 that can prevent it since commercial vaccines available in Brazil and worldwide, only protect against EHV-1 and 4. The relation between all the abortions with EHV-2 can’t be confirmed once samples from these animals were not collected. Otherwise, the necropsy of 10 foals indicate similar results with the observations from the foal sampled. All foals died maybe due to this strain bee more pathogenic, since no other agent was isolated and we are not prepared to vaccinated and prevent future EHV-2 outbreaks.

Our study has several limitations including (i) sampling of only one foal fatal case; (ii) absence of pulmonary tissue for molecular detection of EHV-2 and (iii) the sequencing of only one sample obtained from fatal case in a distinct organ related with the probability cause of death. The strengths of our study include (i) sampling of distinct tissues of a dead foal from a farm presenting consecutive horse abortions and foals death an outbreak with unidentified pathogen or cause, (ii) sensitive methodology targeting major respiratory/neurologic pathogens, (iii) the combination of histopatological, necropsy and virological data.

## 5. Conclusion

We report the potential relation of Equine Herpesvirus Type 2 (EHV-2) with horse disease outbreak of respiratory and neurologic disease in a farm of Brazil leading to the death of 11 foals. This is the first EHV-2 outbreak in Brazil showing disease and leading to dead, confirmed by PCR method. The acute characteristics and high morbidity highlight the importance of this virus circulation. Furthermore, conventional treatment was unsuccessful, maybe due to a higher pathogenicity of this strain, since no other agent was isolated.

This case reports it’s an alert of what can became a huge problem for horse breeders and veterinarians as this virus can spread throw Brazil and South America continent since it’s a border region. Our results confirm the presence of EHV-2 working an important role in equid respiratory disease.

## Highlights

Gammaherpesvirus EHV-2 neglected and circulating in symptomatic equids in South America
EHV-2 outbreak in Middle-west region of Brazil 11 foals dead.
Foals died with respiratory symptoms evolving to neurological signs
EHV-2 outbreak in Middle-west region of Brazil several mares had aborted

## References

1. Azab W, Bedair S, Abdelgawad A, Eschke K, Farag GK, Abdel-raheim A, et al. Detection of equid herpesviruses among different Arabian horse populations in Egypt. Vet Med Sci. 2019;5:361–71.

2. Kershaw O, Von Oppen T, Glitz F, Deegen E, Ludwig H, Borchers K. Detection of equine herpesvirus type 2 (EHV-2) in horses with keratoconjunctivitis. Virus Res. 2001;80(1–2):93–9.

3. Sledge DG, Miller DL, Styer EL, Hydrick HA, Baldwin CA. Equine herpesvirus 2-associated granulomatous dermatitis in a horse. Vet Pathol. 2006;43(4):548–52.

4. Schlocker N, Gerber-Bretscher R, von Fellenberg R. Equine herpesvirus 2 in pulmonary macrophages of horses. Am J Vet Res. 1995;Jun; 56(6):749–54.

5. Harden TJ, Bagust TJ, Pascoe RR, Spradbrow PB. STUDIES ON EQUINE HERPESVIRUSES 5. Isolation and Characterisation of Slowly Cytopathic Equine Herpesviruses in Queensland. Aust Vet J. 1974;50(11):483–8.

6. Galosi CM, De La Paz VC, Fernández LC, Martinez JP, Craig MI, Barrandeguy M, et al. Isolation of equine herpesvirus-2 from the lung of an aborted fetus. J Vet Diagnostic Investig. 2005;17(5):500–2.

7. Belák S, Pálfi V, Tuboly S, Bartha L. Passive immunization of foals to prevent respiratory disease caused by equine herpesvirus type 2. Zentralblatt fur Vet R B J Vet Med Ser B. 1980;27(9–10):826–30.

8. Pálfi V, Belák S, Molnár T. Isolation of equine herpesvirus type 2 from foals, showing respiratory symptoms. Zentralblatt fur Vet R B J Vet Med Ser B. 1978;25(2):165–7.

9. Giguère S, Prescott JF. Clinical manifestations, diagnosis, treatment, and prevention of Rhodococcus equi infections in foals. Vet Microbiol. 1997;56(3–4):313–34.

10. Giguère S, Cohen ND, Keith Chaffin M, Hines SA, Hondalus MK, Prescott JF, et al. Rhodococcus equi: Clinical Manifestations, Virulence, and Immunity. J Vet Intern Med. 2011;25(6):1221–30.

11. Craig MI, Barrandeguy ME, Fernández FM. Equine herpesvirus 2 (EHV-2) infection in thoroughbred horses in Argentina. BMC Vet Res. 2005;1:1–5.

12. Dall Agnol AM, Beuttemmuller EA, Pilz D, Leme RA, Saporiti V, Headley SA, et al. Detection of Equid gammaherpesvirus 2 and 5 DNA in the upper respiratory tract of asymptomatic horses from Southern Brazil. Brazilian J Microbiol. 2019;2–5 .

13. Negussie H, Gizaw D, Tesfaw L, Li Y, Oguma K, Sentsui H, et al. Detection of Equine Herpesvirus (EHV) −1, −2, −4 and −5 in Ethiopian Equids with and without Respiratory Problems and Genetic Characterization of EHV-2 and EHV-5 Strains. 2017;64:1970–8.

14. Vandevanter DR, Warrener P, Bennett L, Schultz ER, Coulter S, Garber RL, et al. Detection and analysis of diverse herpesviral species by consensus primer PCR. J Clin Microbiol. 1996;34(7):1666–71.

15. Johnson N, Wakeley PR, Mansfield KL, McCracken F, Haxton B, Phipps LP, et al. Assessment of a novel real-time pan-flavivirus RT-polymerase chain reaction. Vector-Borne Zoonotic Dis. 2010;10(7):665–71.

16. Sánchez-Seco MP, Rosario D, Domingo C, Hernández L, Valdés K, Guzmán MG, et al. Generic RT-nested-PCR for detection of flaviviruses using degenerated primers and internal control followed by sequencing for specific identification. J Virol Methods. 2005;126(1– 2):101–9.

17. Marenzoni ML, Bietta A, Lepri E, Casagrande Proietti P, Cordioli P, Canelli E, et al. Role of equine herpesviruses as co-infecting agents in cases of abortion, placental disease and neonatal foal mortality. Vet Res Commun. 2013;37(4):311–7.

18. Léon A, Fortier G, Fortier C, Freymuth F, Tapprest J, Leclercq R, et al. Detection of equine herpesviruses in aborted foetuses by consensus PCR. Vet Microbiol. 2008;126(1–3):20–9.

19. Marenzoni M, Stefanetti V, Danzetta ML, Timoney PJ. Gammaherpesvirus infections in equids: a review. Vet Med Res Reports [Internet]. 2015 Apr 1 [cited 2019 Jul 31];6:91. Available from: http://www.dovepress.com/gammaherpesvirus-infections-in-equids-a-review-peer-reviewed-article-VMRR

20. Woźniakowski G, Samorek-Salamonowicz E. Animal herpesviruses and their zoonotic potential for cross-species infection. Ann Agric Environ Med. 2015;22(2):191–4.

21. Garcia Ribeiro M, Seki I, Yasuoka K, Kakuda T, Sasaki Y, Tsubaki S, et al. Molecular epidemiology of virulent Rhodococcus equi from foals in Brazil: Virulence plasmids of 85-kb type I, 87-kb type I, and a new variant, 87-kb type III. Comp Immunol Microbiol Infect Dis. 2005;28(1):53–61.

22. Patel JR, Bateman H, Williams J, Didlick S. Derivation and characterisation of a live equid herpes virus-1 (EHV-1) vaccine to protect against abortion and respiratory disease due to EHV-1. Vet Microbiol. 2003;91(1):23–39.

